# Initial HCV infection of adult hepatocytes triggers a temporally structured transcriptional program containing diverse pro- and anti-viral elements

**DOI:** 10.1101/2021.02.12.431054

**Authors:** Birthe Tegtmeyer, Gabrielle Vieyres, Daniel Todt, Chris Lauber, Corinne Ginkel, Michael Engelmann, Maike Herrmann, Christian K. Pfaller, Florian W. R. Vondran, Ruth Broering, Ehsan Vafadarnejad, Antoine-Emmanuel Saliba, Christina Puff, Wolfgang Baumgärtner, Csaba Miskey, Zoltán Ivics, Eike Steinmann, Thomas Pietschmann, Richard J. P. Brown

**Affiliations:** Institute of Experimental Virology, TWINCORE, Centre for Experimental and Clinical Infection Research; a joint venture between the Medical School Hannover (MHH) and the Helmholtz Centre for Infection Research (HZI), Hannover, Germany; Heinrich Pette Institute, Leibniz Institute for Experimental Virology, Hamburg, Germany; Department of Molecular and Medical Virology, Bochum, Germany; European Virus Bioinformatics Center (EVBC), Jena, Germany; Division of Veterinary Medicine, Paul Ehrlich Institute, Langen, Germany; Department of General, Visceral and Transplant Surgery, Hannover Medical School, 30625 Hannover, Germany; German Centre for Infection Research (DZIF), partner site Hannover-Braunschweig, Hannover, Germany; Department of Gastroenterology and Hepatology, University Hospital Essen, University Duisburg-Essen, Essen, Germany; Helmholtz Institute for RNA-based Infection Research (HIRI), Helmholtz-Center for Infection Research (HZI), Würzburg, Germany; Department of Pathology, University of Veterinary Medicine Hannover, Hannover, Germany; Division of Medical Biotechnology, Paul Ehrlich Institute, Langen, Germany

**Keywords:** Hepatitis C virus (HCV), RNA-seq, primary human hepatocytes, IFN regulatory factor 1 (*IRF1*), IFN signaling, EIF2 signaling, translational shut-off

## Abstract

Transcriptional profiling provides global snapshots of virus-mediated cellular reprogramming, which can simultaneously encompass pro- and antiviral components. To determine early transcriptional signatures associated with HCV infection of authentic target cells, we performed ex vivo infections of adult primary human hepatocytes (PHHs) from seven donors. Longitudinal sampling identified minimal gene dysregulation at six hours post infection (hpi). In contrast, at 72 hpi, massive increases in the breadth and magnitude of HCV-induced gene dysregulation were apparent, affecting gene classes associated with diverse biological processes. Comparison with HCV-induced transcriptional dysregulation in Huh-7.5 cells identified limited overlap between the two systems. Of note, in PHHs, HCV infection initiated broad upregulation of canonical interferon (IFN)-mediated defense programs, limiting viral RNA replication and abrogating virion release. We further find that constitutive expression of *IRF1* in PHHs maintains a steady-state antiviral program in the absence of infection, which can additionally reduce HCV RNA translation and replication. We also detected infection-induced downregulation of ∼90 genes encoding components of the EIF2 translation initiation complex and ribosomal subunits in PHHs, consistent with a signature of translational shutoff. As HCV polyprotein translation occurs independently of the EIF2 complex, this process is likely pro-viral: only translation initiation of host transcripts is arrested. The combination of antiviral intrinsic and inducible immunity, balanced against pro-viral programs, including translational arrest, maintains HCV replication at a low-level in PHHs. This may ultimately keep HCV under the radar of extra-hepatocyte immune surveillance while initial infection is established, promoting tolerance, preventing clearance and facilitating progression to chronicity.

**IMPORTANCE:** Acute HCV infections are often asymptomatic and therefore frequently undiagnosed. We endeavored to recreate this understudied phase of HCV infection using explanted PHHs and monitored host responses to initial infection. We detected temporally distinct virus-induced perturbations in the transcriptional landscape, which were initially narrow but massively amplified in breadth and magnitude over time. At 72 hpi, we detected dysregulation of diverse gene programs, concurrently promoting both virus clearance and virus persistence. On the one hand, baseline expression of *IRF1* combined with infection-induced upregulation of IFN-mediated effector genes suppresses virus propagation. On the other, we detect transcriptional signatures of host translational inhibition, which likely reduces processing of IFN-regulated gene transcripts and facilitates virus survival. Together, our data provide important insights into constitutive and virus-induced transcriptional programs in PHHs, and identifies simultaneous antagonistic dysregulation of pro-and anti-viral programs which may facilitate host tolerance and promote viral persistence.

## INTRODUCTION

Despite development of effective antiviral therapies, hepatitis C virus (HCV) remains a global health burden and still chronically infects around 71 million people worldwide. HCV is a positive-stranded RNA virus of the *Flaviviridae* family with a tropism restricted to human hepatocytes (1). The development of cell culture systems has yielded insights into HCV-host interactions. For instance, a lipid metabolic reprogramming favorable to the constitution of the viral replication organelle (2) and stress responses (3) were reported upon HCV infection. These dysregulations facilitate viral propagation but also the development of pathogenesis in chronic infection, including liver steatosis (4). Balanced against this, HCV infection elicits a classical innate immune response that obeys rules that are conserved across viruses (5). Virus infection is sensed via diverse pattern recognition receptors (PRRs) which trigger signaling cascades involving the nuclear translocation of IFN regulatory factors (IRFs) and NF-κB (nuclear factor-κB). These transcription factors activate the expression of target genes, including IFNs, which in turn activate the transcription of a panel of IFN-regulated genes (IRGs) in a paracrine and autocrine manner (6). Many of these IRGs have known antiviral effects (7). Importantly, these host protective responses are dampened by the virus, for example by the viral protease NS3-4A, which cleaves several key molecules in the IFN induction pathway, including the adaptor MAVS (5).

Many of these changes, whether metabolic reprogramming to facilitate viral replication or defense reactions, are accompanied or caused by transcriptional alterations of the infected cell (8, 9). Transcriptome-wide studies of HCV infection mostly focused so far on cell lines, in particular on the Huh-7.5 hepatoma cell line (10), which represents a robust *in vitro* infection model (e.g. (8, 9)). Transcriptional profiling of liver biopsies from chronically infected patients were also performed (11). Previous studies in primary human hepatocytes (PHHs) were based on RT-qPCR or microarrays and restricted to specific gene subsets, in particular focusing on innate immunity (12–14). Responses to HCV infection in infected human hepatocytes at near-single cell resolution were described. However these cells were derived from fetal liver cells rather than directly explanted from an adult liver (15) and have an immature phenotype (16). Additionally, the initial phase of infection directly preceding infection was not captured.

In this report, we used RNA-seq (17) to analyze the host transcriptional landscape directly after HCV infection of adult PHHs. Infection of human adult hepatocytes plated from liver resections of HCV-negative patients opens the unique possibility to study early time points directly after infection of natural HCV target cells, without contamination from other liver cell types (18) which could blur transcriptional signals. We compared the responses at 6 hpi and 72 hpi in order to reflect the spatial transitioning of viral replication complexes from ribosomes (19) to endoplasmic reticulum (20). In parallel, we also performed transcriptional profiling of highly permissive Huh-7.5 cells, which are widely used to propagate HCV *in vitro*, and directly compare HCV-induced gene dysregulation in the two systems for the first time.

## RESULTS

### Experimental design and data visualization

Explanted adult PHHs from seven donors (D1-7) were plated and checked for viability prior to infection experiments. PHHs were incubated with conditioned medium (CM) or infected with replication competent HCV (strain Jc1). Additionally, we inoculated PHHs with UV-inactivated HCV where sufficient viable patient material was available (HCV^UV^: D4-7). Cellular RNAs were isolated at two early time points (6 hpi and 72 hpi) to enable monitoring of global hepatocyte transcriptional changes by RNA-seq and quantification of HCV RNA (Fig 1A). Infection experiments in Huh-7.5 cells were also performed in parallel.

**Figure 1:**
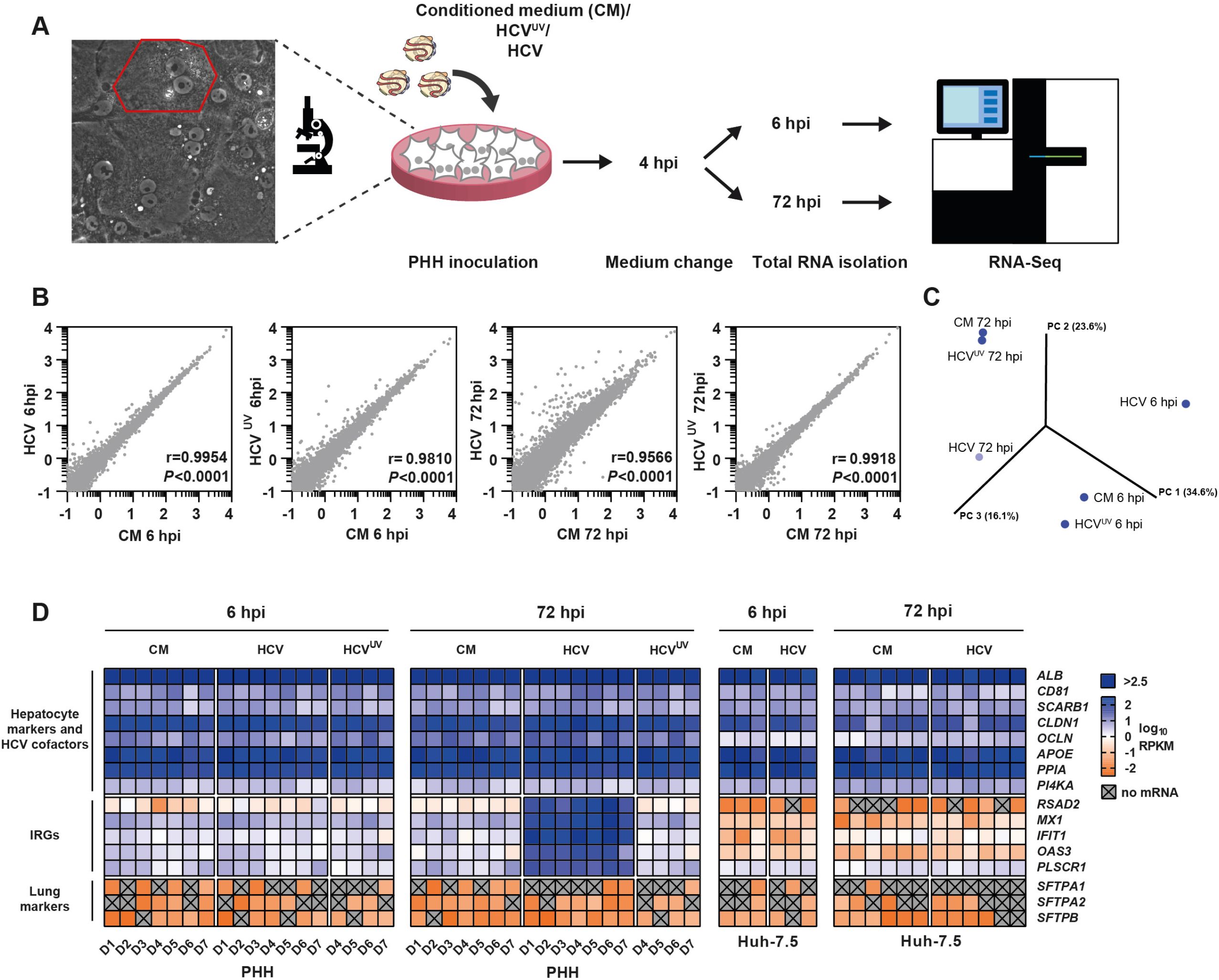
Experimental protocol and RNA-seq data validation. **(A)** Schematic of experimental protocol. PHH: primary human hepatocytes. RNA-seq: RNA sequencing. **(B)** Visualization of global transcriptional changes induced by HCV infection (n=7 donors) or treatment with UV-inactivated HCV (HCV^UV^, n=4 donors). For individual plots, average RPKM (log_10_) values for all detected transcripts from conditioned medium (CM) treated cells are plotted on the x-axes, with corresponding values from HCV-infected and HCV^UV^ treated cells plotted on the y-axis, respectively. Pearson’s r and p-values for each comparison are inset. hpi: hours post infection. **(C)** Three dimensional principal component analysis (PCA) from PHH donor 4. Samples are distinguishable depending on their time point and infection status. **(D)** Control gene expression levels in PHHs and Huh-7-5 cells at 6 and 72 hpi. Heat maps of control gene RPKM values (log_10_) from individual PHH donors or different Huh-7.5 cell passages. IRGs: Interferon regulated genes.

After mapping RNA-seq data to the hg38 genome scaffold, raw count data were normalized (reads per kilobase of transcript per million mapped reads: RPKM) to allow comparison of global gene expression profiles within and between experiments. Mean RPKM values for all expressed genes were plotted for CM-treated PHHs versus HCV or HCV^UV^ infected PHHs, and correlation analyses performed at both 6 hpi and 72 hpi (Fig 1B). These analyses revealed all comparisons were highly significant (*P*<0.0001) with Pearson’s r correlation close to 1, indicating the majority of hepatocyte mRNAs are expressed at steady-state and not significantly dysregulated upon HCV infection. Of note, Pearson’s r was lowest at 72 hpi with replication competent HCV (0.9566), indicating greater numbers of dysregulated hepatocyte genes under these conditions (Fig 1B).

Principle component analyses (PCA), performed separately on samples from individual donors, revealed HCV-infected PHHs were clearly separated from CM-treated and HCV^UV^ infected cells at both time points (see example plot from D4, Fig 1C). In addition, for all donors, temporally distinct clusters at 6 and 72 hours were apparent, validating our approach of using time-matched infected and uninfected PHHs to avoid mixing of signals associated with HCV infection versus gradual hepatocyte de-differentiation upon plating.

### HCV infection activates antiviral defenses in PHHs but not Huh-7.5 cells

Transcript abundance of a panel of selected control genes were compared in PHHs and highly permissive Huh-7.5 cells (21), both with and without HCV infection. Comparable abundant expression of hepatocyte marker *ALB* and transcripts encoding HCV entry and replication co-factors were detected in both PHHs and Huh-7.5 cells. Their expression remained stable across the experimental time-course suggesting no reduction in HCV permissiveness and no modulation of expression due to infection (Fig 1D). As expected, expression of lung-specific transcripts was either minimal or absent.

Baseline expression of a panel of IFN regulated genes (IRGs) was detectable in PHHs, which was not further boosted upon infection with HCV^UV^ at either 6 or 72 hpi. While minimal IRG upregulation was observed upon infection with replication competent HCV at 6 hours, substantial induction was observed at 72 hpi. In contrast, basal IRG expression was demonstrably lower or completely absent in Huh-7.5 cells when compared to PHHs, and no induction was observed upon HCV infection at either sampling point.

### Antiviral defenses in PHHs suppress HCV replication and completely abrogate virion release

To validate our RNA-seq data, we performed RT-qPCR on a selected set of test genes and compared levels of gene induction in both systems. These data confirm a remarkable level of concordance between the two systems and confirm our RNA-seq data accurately records both steady-state and virus-inducible gene expression (Fig 2A). To determine HCV infection rates and investigate the effects of IRG induction on HCV RNA replication, vRNA RT-qPCR was performed on cellular RNAs from PHHs and Huh-7.5 cells. Individual cells contain between 10-30pg total RNA: 1µg therefore represents 3.3×10^4^ -1.0×10^5^ cells. At 6 hpi in PHHs we observed a mean of 6.2×10^4^ HCV GE/ug total RNA, which equates to an initial infection rate of 0.5 – 1.6 viral genome copies per cell (Fig 2B, top panel). This copy number remained stable at 72 hpi, likely reflecting low-level HCV replication with suppression mediated by PAMP-induced innate immunity. In contrast, while slightly lower infection rates were observed in Huh-7.5 cells at 6 hpi, a 2-log increase in vRNA was apparent at 72 hpi (Fig 2B, bottom panel), consistent with a lack of IRG upregulation in Huh-7.5 cells. As expected, minimal signal was detected in CM or HCV^UV^ treated cells. To determine rates of infectious particle production, TCID_50_ titrations were performed in parallel on supernatants harvested from infected PHHs and Huh-7.5 cells (Fig 2C). Titers at 6 hpi likely represent carryover of initial inoculum, despite extensive washing. While infectious virion secretion was completely absent in supernatants from PHHs at 72 hpi, HCV virions were detected at ∼1×10^5^ TCID_50_/ml in supernatants from Huh-7.5 cells at 72 hpi. To investigate this further, virion secretion was determined in PHHs which were pre-treated with the JAK/STAT inhibitor ruxolitinib prior to infection. Ruxolitinib pre-treatment rescued HCV virion release at 72 hpi indicating the ablation of virion production in PHHs is mediated by JAK/STAT-inducible immunity (Fig 2C, top panel). Moreover, these results were confirmed via immunofluorescence staining for viral antigen. At 72 hpi, perinuclear NS5A localization in PHHs was only visible after JAK/STAT inhibition (Fig 2D) but readily detected in Huh-7.5 cells without pharmacological immune suppression (Fig 2E). Together, these data indicate that plated PHHs possess intact innate immunity which suppresses initial HCV replication and completely blocks virion release. This inducible immunity is absent in Huh-7.5 cells.

**Figure 2:**
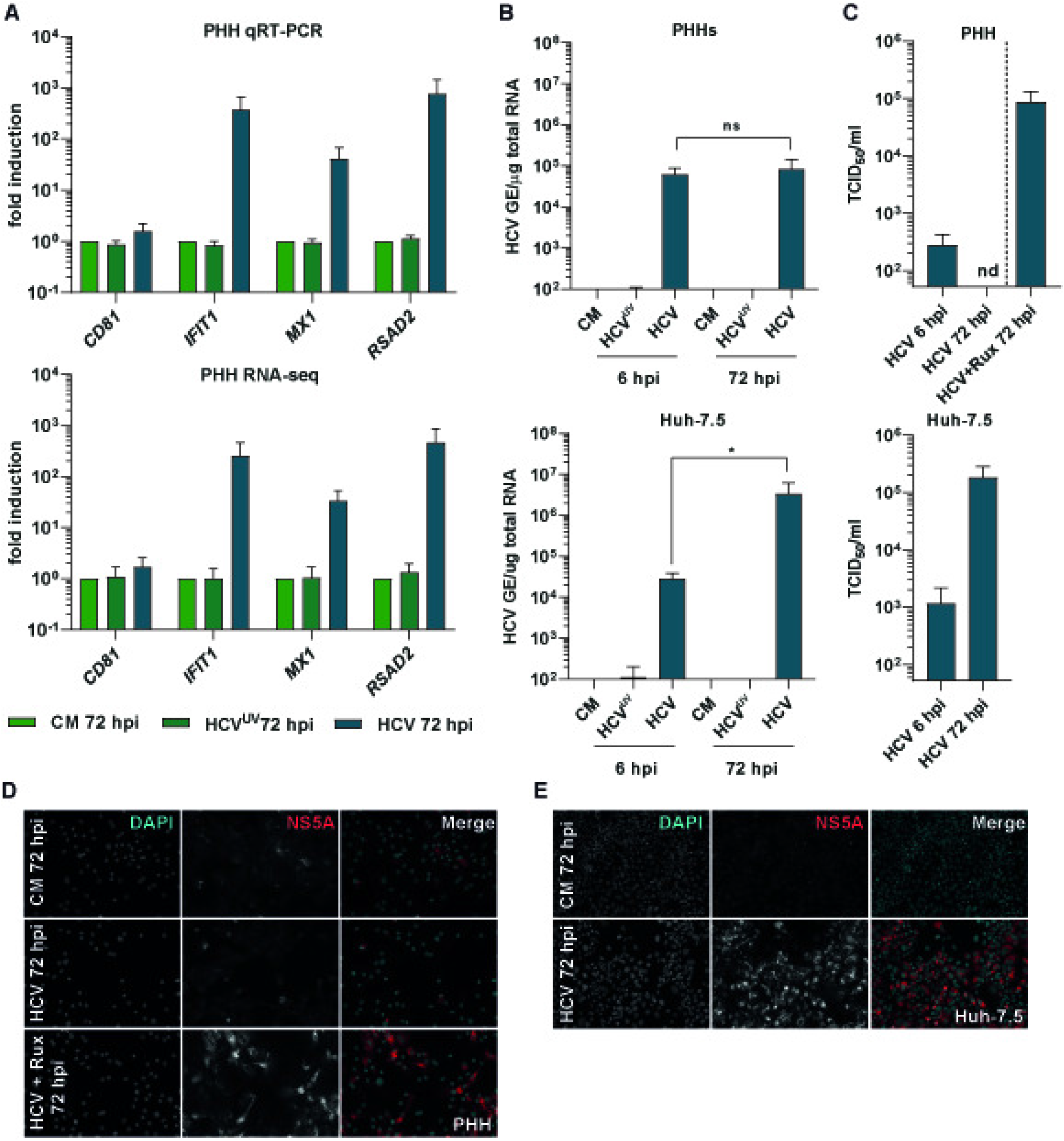
Intact innate immunity suppresses HCV propagation in PHHs. **(A)** Comparative gene induction of four control genes measured by RT-qPCR or RNA-seq (n=4 donors). Top panel shows fold gene induction determined by RT-qPCR and calculated by the 2^-ΔΔCT^ method (43) compared to CM treated PHHs. Bottom panel shows fold gene induction based on RNA-seq data for the same samples under identical conditions. **(B)** Intracellular HCV-RNA copies in HCV infected PHHs (n=4 donors, top) and Huh-7.5 cells (n=3, bottom). Bars represent HCV RNA GE (genome equivalents) per 1µg total RNA. *: p-value < 0.05. ns: not significant. Data was log-transformed and a one-tailed, unpaired t-test applied to each comparison. **(C)** Secretion of infectious virions in supernatants of HCV infected PHHs (top) and Huh-7.5 cells (bottom). Numbers of infectious particles in PHH supernatents at 72 hpi were under the LOQ of the assay at a 1:3 dilution. Dashed line indicates a separate experiment with four additional PHH donors preincubated with 10µM ruxolitinib (Rux), a JAK/STAT inhibitor, prior to and directly after HCV infection. **(D-E)** Immunofluorescence staining of viral NS5A protein in PHH **(D)** or Huh-7.5 cells. **(E)** PHH were treated with CM or infected with HCV for 72 hours with or without ruxolitinib pretreatment. NS5A staining is shown in red with nuclear counterstaining (DAPI) in blue.

### HCV infection of PHHs dysregulates gene programs associated with diverse biological functions

Statistical analyses were performed to quantify differentially expressed genes (DEGs) induced upon HCV infection of PHHs (FDR *P*<0.05). For replication competent HCV infections, a temporally structured increase in DEGs was observed (Fig 3A, left panel and Fig 3B) which partially overlapped (Fig. 3C). At 6 hpi, n=81 infection-induced DEGs were apparent – this number was markedly amplified at 72 hpi (DEGs n=2985) and indicates HCV-infection ultimately induces a substantial shift in the PHH transcriptional landscape (Fig 3A, left panel and 3B). For HCV^UV^ infections, a contrasting temporally-structured reduction in DEG signatures was observed: an initial transcriptional response to HCV^UV^ was detected at 6 hpi (n=239) which declined to virtually no detectable dysregulation at 72 hpi (n=12) (Fig 3A, left panel).

**Figure 3:**
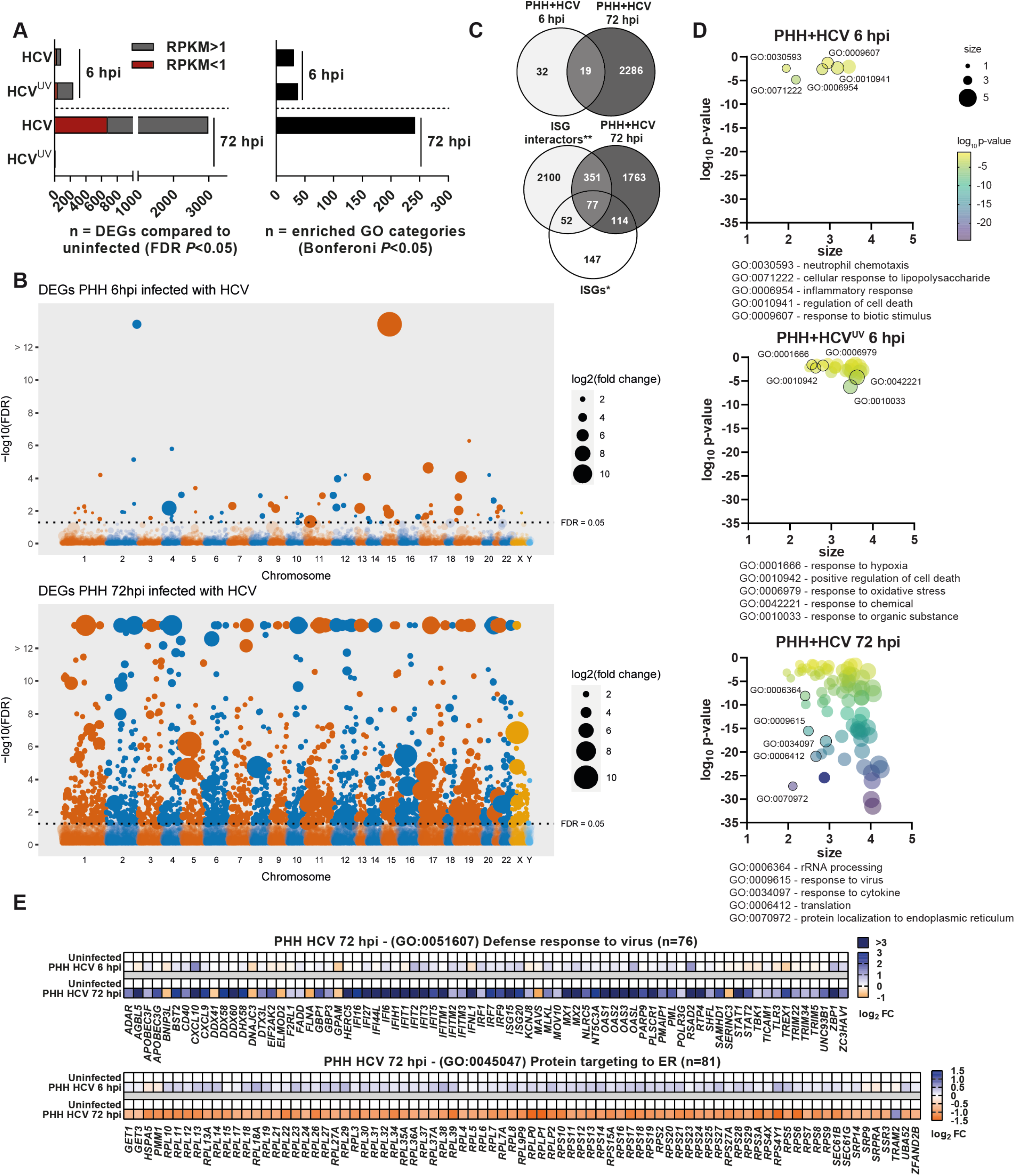
HCV infection induces temporally structured and functionally diverse gene programs in PHHs. **(A)** Comparison of numbers of HCV and HCV^UV^ induced DEGs (left) and enriched GO categories (right) at 6 and 72 hpi. **(B)** Genomic location of HCV-induced transcripts. Manhattan plots compare transcript abundance in uninfected versus HCV-infected PHHs at 6 hpi (top) and 72 hpi (bottom). Circles represent individual gene comparisons with sizes proportional to average RPKM fold-change (log_2_). **(C)** Number of HCV-induced genes in PHH at 6 and 72 hpi. Top Venn diagram shows overlap between DEGs at 6 and 72 hpi, while gene overlap at 72 hpi with characterized IRGs* (7) and IRG interactors** (22) are displayed below. Only DEGs with FDR p-value <0.05 and RPKM >1 were included. **(D)** GO enrichment analysis of HCV-induced DEGs in PHHs. Each circle represents a GO category and is plotted depending on its p-value and size. Size illustrates the frequency of the GO term in the underlying GO annotation (46). Representative GO categories are highlighted in graphs and their full annotation displayed below each plot. **(E)** HCV-induced transcripts at 72 hpi. Mean (n=7 donors) fold change (log_2_) of HCV-induced DEGs associated with defense response to virus (top) or protein targeting to ER (bottom).

To determine the biological processes associated with HCV-induced DEGs, gene ontology (GO) enrichment analyses were performed. At 6 hpi, HCV and HCV^UV^ enriched GO categories were associated with shared and distinct biological processes (Fig 3A right panel and Fig 3D, two upper panels). Of note, only PHHs infected with replication competent HCV demonstrated enrichment of GO categories associated with IFN signaling or pathogen defense: dysregulation of these gene classes were absent in HCV^UV^ infected PHHs. These data represent the early yet restricted signatures of the hepatocyte antiviral response and confirm this response is induced only by replication competent vRNAs.

Proportional to the quantity of HCV-induced DEGs, a large number of highly significant enriched GO categories were identified at 72 hpi, which were associated with diverse biological processes (Fig 3A, right panel and 3D, bottom panel). Overlap with DEGs representing described IRGs (n=390) (7, 22) or IRG-interactors (n=2582) (22) (Fig 3C, bottom panel) resulted in enrichment of multiple GO categories associated with antiviral responses and innate immunity (Fig 3D, bottom panel). However, many DEGs were not classical IRGs/IRG-interactors and represent biological processes not automatically associated with the antiviral response (eg. ribosome biogenesis, translation initiation and protein targeting to the ER) (Fig 3D, bottom panel). Examples of opposing patterns of gene dysregulation can be seen in Fig 3E for two representative GO categories: while genes associated with defense response to virus are generally strongly upregulated at 72 hpi, genes associated with protein targeting to the ER are almost exclusively down regulated. Together, these early snap-shots of virus-induced cellular changes confirm temporally regulated transcriptional responses to HCV infection in adult PHHs, with the magnitude of gene induction increasing exponentially from 6 to 72 hpi.

### Divergent transcriptional responses to HCV infection and ectopic *IRF1* expression in Huh-7.5 cells

Huh-7.5 cells represent the most frequently used cell-line for HCV propagation in research. Consequently, we also cataloged global HCV-induced transcriptional responses in these cells for comparative purposes. Similar to PHH, statistical analyses identified a time-dependent increase in significant HCV-induced DEGs and enriched GO categories (Fig 4A).

**Figure 4:**
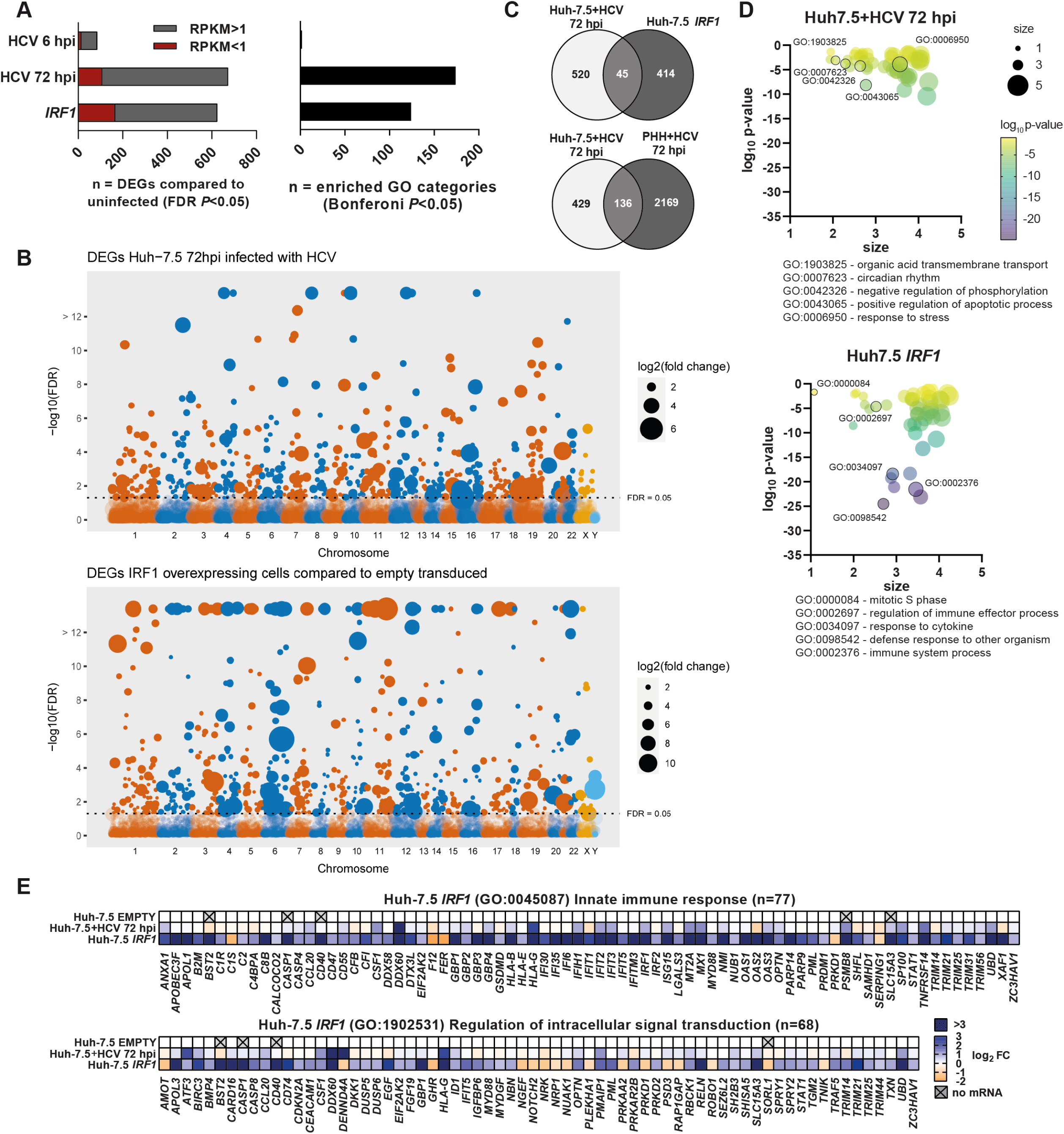
Distinct *IRF1*- and HCV-mediated transcriptional programs in Huh-7.5 cells. **(A)** Comparison of numbers DEGs (left) and enriched GO categories (right) upon HCV infection or ectopic *IRF1*-expression in Huh-7.5 cells. **(B)** Genomic location of HCV-induced (top) or *IRF1*-induced transcripts (bottom) in Huh-7.5 cells. Manhattan plots compare transcript abundance in uninfected Huh7-5 cells to HCV-infected or *IRF1*-expressing cells. Circles represent individual gene comparisons with sizes proportional to average RPKM fold-change (log_2_). **(C)** HCV-and *IRF1*-induced DEGs show limited overlap. Top Venn diagram show overlap between HCV-and *IRF1*-induced DEGs while overlap of HCV-induced DEGs in Huh-7.5 versus PHHs is shown below. Only DEGs with FDR p-value <0.05 and RPKM >1 were included. **(D)** GO enrichment analyses of HCV-and *IRF1*-induced DEGs in Huh-7.5 cells. Each circle represents a GO category and is plotted depending on its p-value and size. Size illustrates the frequency of the GO term in the underlying GO annotation (46). Representaive GO categories are highlighted in graphs and their full annotation displayed below each plot. **(E)** *IRF1*-induced transcripts in Huh-7.5. Mean fold change (log_2_) of *IRF1*-induced DEGs associated with innate immune response (top) or regulation of intracellular signal transduction (bottom).

Intrinsic *IRF1* expression has been reported to maintain the baseline transcription of a program of antiviral genes, independently of the IFN system (23, 24). To quantify the *IRF1*-regulon in human cells of hepatic origin, we transcriptionally re-programmed Huh-7.5 cells by ectopically expressing *IRF1*. DEGs and GO enriched categories induced by HCV at 72 hpi, and ectopic *IRF1* expression were numerically similar (Fig 4A and 4B). However, the underlying dysregulated genes exhibited approximately 10% overlap (Fig 4C, upper panel) and the significantly enriched biological processes associated with the dysregulated genes were highly divergent (Fig 4D). HCV induced genes in Huh-7.5 cells were associated with diverse biological processes presumably facilitating viral propagation, including circadian rhythm or amino acid transport (Fig 4D, upper panel) and showed minimal overlap with HCV-induced genes in PHHs (Fig 4C, lower panel). In contrast, *IRF1* regulated genes were largely but not exclusively associated with GO categories related to innate immunity or pathogen defense (Fig 4D, lower panel). Examples of *IRF1*-mediated gene dysregulation are shown for two representative GO categories associated with innate immunity (Fig 4E). In summary, these data expand the *IRF1*-regulon identified by gene microarrays in (7) (n=130) by an additional 329 genes (n=459) and confirm that Huh-7.5 cells retain the capacity to mount antiviral defenses. Our highly sensitive RNA-seq profiling identified an additional 271 low-abundance transcripts as significantly dysregulated (FDR p-values <0.05, final RPKM <1) although these were omitted from subsequent analyses to increase stringency.

### Baseline *IRF1* expression coordinates an intrinsic antiviral program in PHHs

Using our previously selected panel of control genes (Fig 1D), we visualized baseline expression of IRGs in uninfected Huh-7.5 cells transduced with an EMPTY lentivirus, Huh-7.5 *IRF1* cells and uninfected PHHs. We observed highly similar baseline IRG expression in Huh-7.5 *IRF1* cells and uninfected PHHs, distinct from Huh-7.5 EMPTY cells (Fig 5A, upper panel). We also visualized baseline expression of *IRF1* in Huh-7.5 cells and PHHs, and compared expression levels to pattern recognition receptors (PRRs) which recognize cytosolic dsRNA (Fig 5A, lower panel). These data confirm minimal *IRF1* expression in Huh-7.5 cells, 4-fold lower than PHH baseline expression. These observations further reveal much higher baseline expression of cytosolic dsRNA sensors *TLR3, DDX58* (RIG-I) and *IFIHI* (MDA5) in PHHs when compared to Huh-7.5 cells, which likely contribute to the differences observed in inducible antiviral immunity.

**Figure 5:**
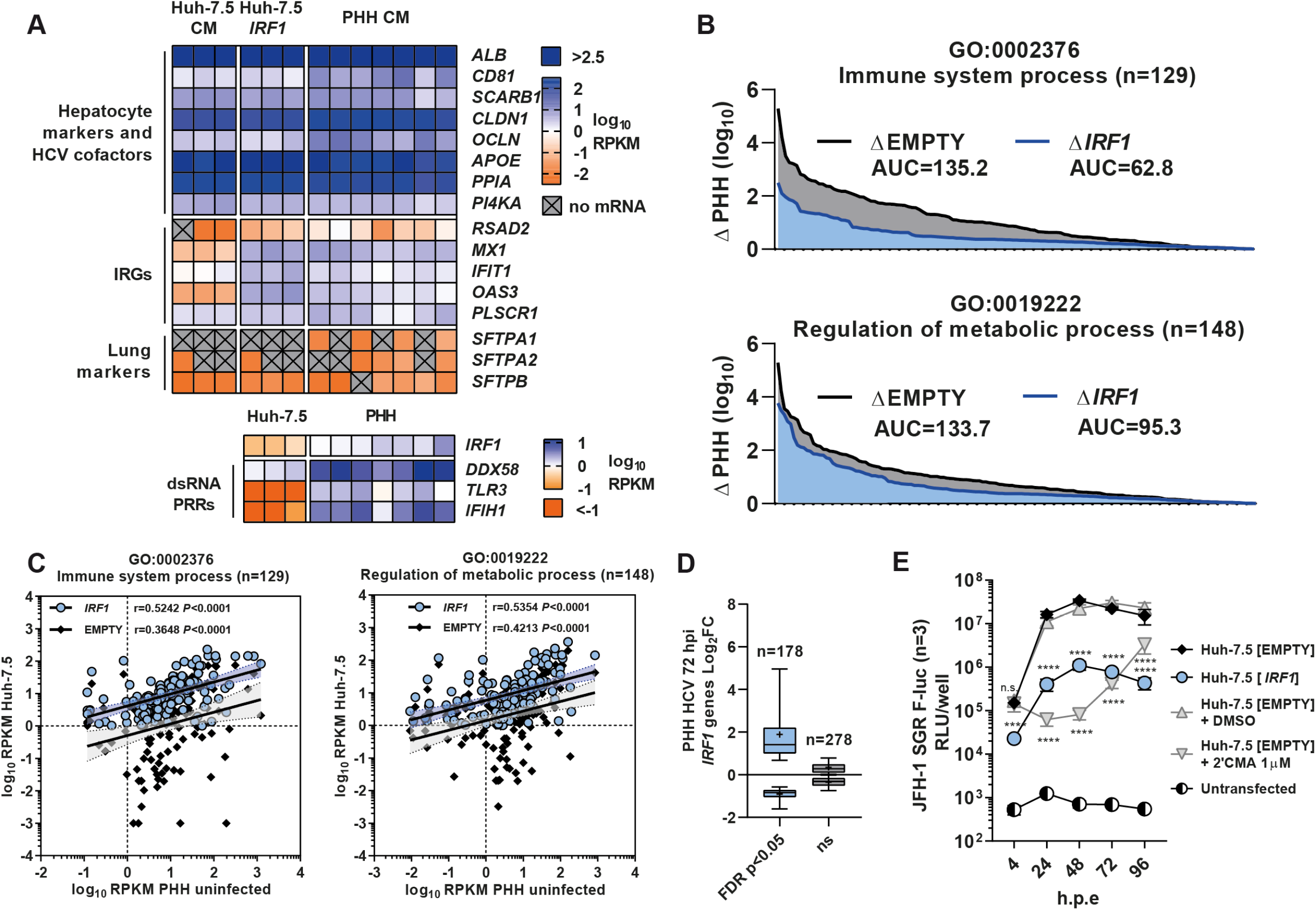
Ectopic *IRF1* expression induces an antiviral gene signature in Huh-7.5 cells similar to baseline expression in PHHs. **(A)** Baseline expression of IRGs and *IRF1*. Top panel. Basal IRG expression in Huh-7.5 with and without *IRF1*-expression, compared to uninfected PHHs. Bottom panel. Comparison of baseline expression of *IRF1* and pattern recognition receptors (PRRs) that recognise double stranded (ds) RNA in Huh-7.5 cells and PHHs. **(B)** Area under the curve (AUC) analysis. Differences in *IRF1*-regulated gene expression levels for two distinct GO categories from Huh-7.5 [EMPTY] and Huh-7.5 [*IRF1*] cells, compared to expression in PHHs. Greater AUC values indicate more divergent expression profiles. **(C)** Correlation plots of *IRF1* regulated genes from the same two GO categories. RPKM values are plotted for individual genes, and simultaneous comparison of their expression in uninfected PHH to both Huh-7.5 [EMPTY] and Huh-7.5 [*IRF1*] cells is visualized. Pearson’s r and p-values for each comparison are inset. **(D)** HCV-inducible *IRF1*-regulated genes in PHHs at 72 hpi. +: mean log_2_ fold change in expression **(E)** Restriction of HCV replication and translation by ectopic *IRF1*-expression. Huh-7.5 [EMPTY] and Huh-7.5 [*IRF1*] cells were electroporated with a subgenomic replicon of JFH-1. h.p.e.: hours post electroporation. ****: p-value < 0.0001. Data was log-transformed and multiple unpaired t-tests were performed comparing Huh-7.5 [EMPTY] vs. Huh-7.5 [IRF1] and Huh-7.5 [EMPTY] + DMSO vs. Huh-7.5 [EMPTY] + 2’CMA, respectively.

Next, we compared mean expression of *IRF1* regulated genes from both Huh-7.5 EMPTY and Huh-7.5 *IRF1* cells to expression in PHHs. Applying robust statistical methods to compare gene subsets from two distinct enriched GO categories within the *IRF1*-regulon, we performed area under the curve (AUC) and correlation analyses. AUC analyses confirmed that expression of *IRF1* regulated genes in Huh-7.5 *IRF1* cells was more closely related to baseline expression in PHH than in Huh-7.5 EMPTY cells (Fig 5B). Of note, this pattern was more pronounced for genes associated with immune system process (GO: 000 2376) than for genes associated with regulation of metabolic process (GO:0019222). Correlation analyses compared mean expression of individual *IRF1* regulated genes in PHHs to both Huh-7.5 EMPTY and Huh-7.5 *IRF1* cells (Fig 5C). While all comparisons were significantly correlated (*P*<0.0001), higher Pearson’s r values were observed for Huh-7.5 *IRF1* cells (r=0.52, and r=0.53) than for Huh-7.5 EMPTY cells (r=0.36, and r=0.42), indicating that ectopic expression of *IRF1* pushes the Huh-7.5 cell transcriptome to a more PHH-like state. Correspondingly, this pattern was again more pronounced for immune system process genes (GO: 0002376) than genes associated with regulation of metabolic process (GO:0019222) (Fig 5C). *IRF1* also represents an IRG and its expression is further boosted by HCV infection (Fig 3E, upper panel). Consequently, we next determined what proportion of the *IRF1*-regulon is further dysregulated upon HCV infection of PHHs (Fig 5D). These analyses identify n=178 *IRF1*-regulated genes who’s expression is significantly dysregulated by HCV infection of PHHs, indicating IRF1 also contributes to inducible immunity.

To determine the effect of ectopic *IRF1* expression on HCV RNA replication, we transfected either Huh-7.5 EMPTY or Huh-7.5 *IRF1* cells with a subgenomic replicon (SGR) containing the non-structural proteins NS3-NS5B from strain JFH-1 coupled to a firefly luciferase (F-luc) reporter, and monitored luciferase accumulation over time (Fig 5E). These experiments confirm that the transcriptional program mediated by *IRF1* has the ability to reduce HCV RNA replication significantly. We also performed transfections in the presence of the HCV replication inhibitor 2’CMA, or a DMSO vehicle control. The significant *IRF1* mediated reduction in RLU was also apparent at 4 hours post transfection in Huh-7.5 *IRF1* cells, which was not the case in 2’CMA treated cells, indicating the *IRF1* gene program also negatively impacts viral genome translation, possibly due to direct targeting or competition with host IRG mRNAs for available ribosomes (Fig 5E). In summary, these analyses provide supportive evidence that *IRF1* coordinates the baseline expression of antiviral effector genes in PHHs in the absence of infection, and also contributes to inducible immunity. The antiviral effect mediated by the *IRF1* gene program was able to reduce HCV replication/translation by 1-2-logs. Thus, in addition to the observed triggering of antiviral defenses by HCV, it is likely that this suite of intrinsically expressed genes in PHHs contributes to defense against incoming virus.

### Comparison of transcriptional regulators and targeted canonical pathways

To determine upstream transcriptional regulators, which orchestrate changes in gene expression induced by HCV infection of PHHs or Huh-7.5 cells, or ectopic *IRF1* expression in Huh-7.5 cells, we performed Ingenuity Upstream Regulator Analysis (Qiagen). Focusing on nuclear transcription factors (TFs), we observed significant activation (blue) or inhibition (orange) (p<0.05) of diverse transcriptional regulators (Fig 6A), with distinct and overlapping TF profiles observed for each dataset. Limited overlap (n=5) was observed between the TFs of coordinating transcriptional responses to HCV infection in PHHs and Huh-7.5 cells. Of note, low oxygen tension has been demonstrated to enhance HCV RNA replication (25) and activation of the hypoxia inducible factor 1-alpha (HIF1A) is observed in both systems, which is a known regulator of transcriptional responses to hypoxia (26).

**Figure 6:**
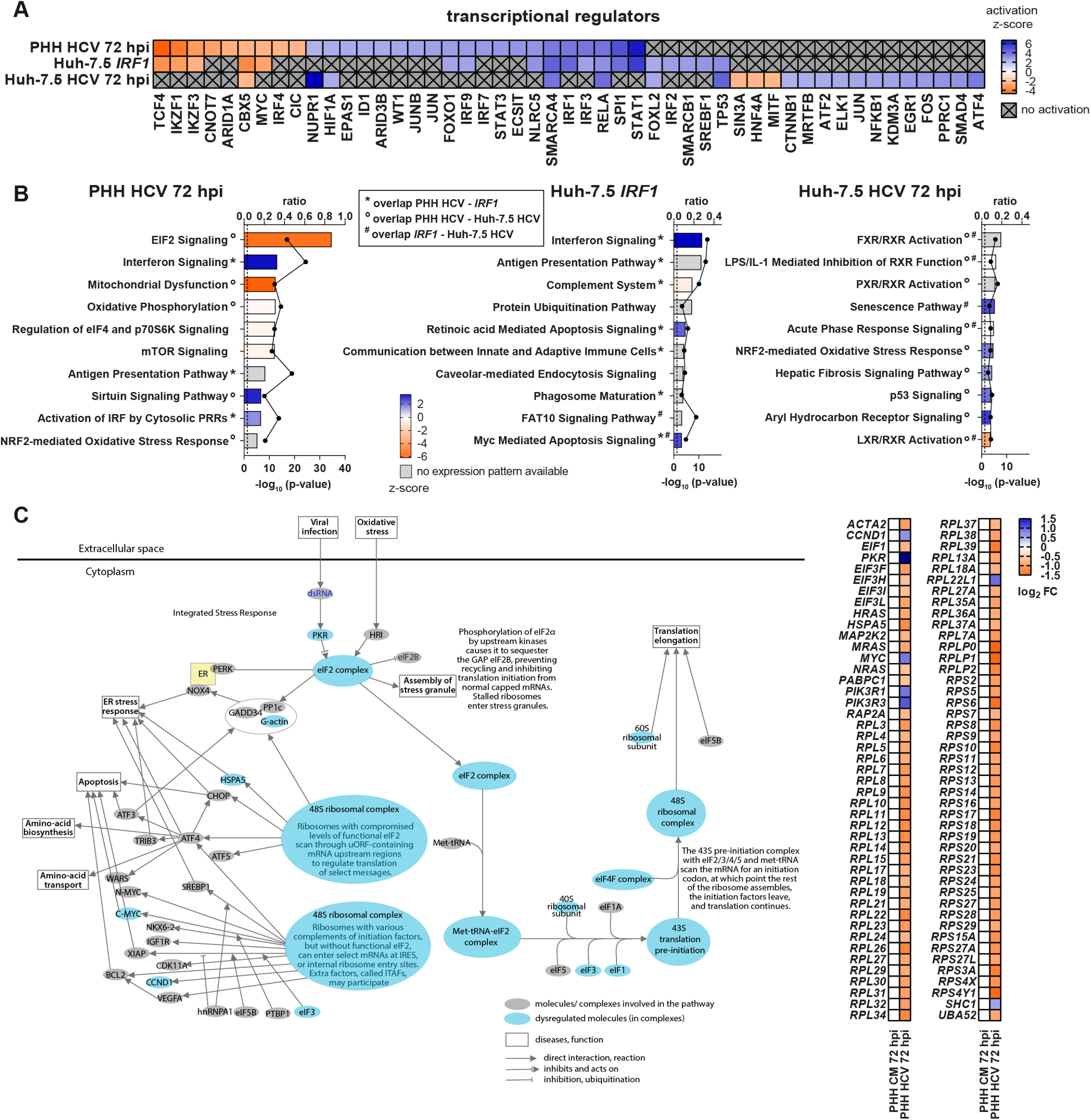
Upstream transcriptional regulators and canonical pathways dysregulated by HCV infection or ectopic *IRF1* expression. **(A)** Upstream transcriptional regulators controlling DEGs. Ingenuity Pathway Analysis (IPA) calculated upstream regulators based on significant DEGs identified under the three presented conditions. Z-scores indicate activation or inhibition of individual regulators. Displayed are only the transcriptional regulators with a p-value <0.05 and a z-score above 2 or lower than -2. **(B)** Canonical pathway analyses of HCV-and *IRF1*-induced DEGs in PHHs and Huh-7.5 cells, respectively. Pathways are plotted with corresponding p-values (bars) and ratios between dysregulated molecules in our datasets and all molecules belonging to that pathway (linked black circles). The dotted line represents the significance threshold (p=0.05). The bar color represents the expression z-score. **(C)** Visualization of the top HCV-dysregulated pathway in PHHs at 72 hpi (EIF2 Signaling) and its underlying DEGs. Left panel shows a modified cartoon of EIF2 Signaling as determined by IPA. Light blue coloring indicates significantly dysregulated molecules associated with the complex/molecule. Heatmap on the right displays fold change of significant DEGs involved in this pathway.

As expected, considerable overlap was observed between the transcriptional regulators coordinating the observed patterns of gene dysregulation detected in HCV infected PHH and *IRF1* reprogramed Huh-7-5 cells, as *IRF1* is further upregulated upon HCV infection of PHHs. Multiple shared TFs known to activate antiviral programs were activated. Furthermore, these analyses demonstrate that ectopic *IRF1* expression additionally activates an array of transcriptional regulators, resulting in a broad induction of antiviral effectors and highlights that *IRF1* co-ordinates a complex web of multiple TFs, which collectively contribute to the *IRF1* regulon.

To further explore these data, we used Ingenuity Pathway Analysis (IPA) (Qiagen) to investigate which canonical cellular pathways are affected by HCV-or *IRF1*-mediated gene dysregulation. Shared and distinct canonical pathways were targeted in all three systems (Fig 6B, only top 10 hits presented). Notably, for HCV infection of PHH, a number of targeted pathways were associated with innate immune responses including ‘Interferon Signaling’, ‘Antigen Presentation Pathway’ and ‘Activation of cytosolic IRF by PRRs’. Targeted transcriptional dysregulation of pathway components associated with IFN-mediated innate immunity were absent in HCV infected Huh-7.5 cells. As Huh-7.5 cells do not produce IFN and possess impaired antiviral effector responses, we reasoned that these HCV-targeted pathways represent pro-viral transcriptional manipulation to facilitate HCV propagation. Cell-intrinsic pathways targeted by *IRF1* were generally involved in innate immunity and exhibited some cross-over with HCV infected PHHs, including ‘Interferon Signaling’, ‘Antigen Presentation Pathway’ and ‘Complement System’.

In HCV infected PHHs, in addition to pathways associated with classical IFN-mediated innate immunity, we also observed targeting of unrelated pathways including ‘EIF2 signaling’, ‘Mitochondrial Dysfunction’ and ‘mTOR signaling’. The top targeted pathway in HCV infected PHHs was ‘EIF2 signaling’ and detailed inspection of this pathway identified significantly dysregulated molecules at multiple pathway stages (Fig. 6B, left panel). Corresponding changes in gene expression for targeted molecules within the ‘EIF2 signaling’ pathway highlight significant downregulation of genes which comprise the structural components of ribosomes and the translation pre-initiation complex. In contrast, significant upregulation *PKR* and *MYC* is observed. (Fig. 6B, right panels). Together these analyses provide a broad overview of the transcriptional regulators which orchestrate HCV or *IRF1*-mediated gene dysregulation, and the downstream cell-intrinsic pathways which they target.

## DISCUSSION

In this study, we sought to quantify and dissect initial global transcriptional responses to HCV infection of authentic target cells -adult PHHs. Acquiring these data *in vivo* is particularly challenging: Indeed, HCV has a highly restricted cellular tropism and efficiently infects only human hepatocytes. The liver is a solid organ which is composed of multiple cell types (27) and while hepatocytes represent the major cell-type, these cells are not readily accessible for sampling. Acute HCV infection is also often asymptomatic and so initial infections often go unnoticed. Here, we sought to overcome these hurdles by performing *ex vivo* infections on PHHs isolated from adult donors that were not previously infected with HCV or treated with IFN. Our quantification of initial acute phase responses to infection in adult PHHs provides a snapshot of early perturbations in the hepatocyte transcriptional landscape induced by HCV infection.

In agreement with reported spatiotemporal shifting of early HCV replication complexes within the cytosol, patterns of HCV-induced gene dysregulation were time-structured in adult PHHs. Limited gene induction was observed at 6 hpi, where initial viral replication complexes are associated with ribosomes (19). However, weak induction of a restricted panel of IRGs was detected, which were not upregulated in HCV^UV^ treated cells, and represents the first signatures of PHHs antiviral response to replication competent HCV. This limited induction of antiviral effectors shortly after infection could represent HCV NS3/4A-mediated targeting of MAVS, which dampens host antiviral responses (28), and contrasts with IFN or PolyI:C treatment of PHHs, where early DEG induction at 6 hours is associated with broad IFN-mediated responses (29). Alternatively, intrinsic *IRF1* expression in PHHs may limit the replication capacity of incoming virus (see below for further discussion), reducing the accumulation of dsRNA and therefore delaying broad innate immune induction. In contrast to 6 hpi DEGs, the spatial transitioning of viral replication complexes to remodeled ER membranes at 72 hpi (20) coincided with an exponential amplification of HCV infection-mediated transcriptional dysregulation. We detected significant dysregulation of ∼3000 genes, associated with a diverse array of biological processes.

At 72 hpi in PHHs, we observed expansive induction of IFN-triggered antiviral effector genes, promoting suppression of viral replication and abrogation of particle release. However, 80% of HCV infected individuals fail to mount effective responses facilitating the progression to chronicity and viruses have evolved a variety of innate immune evasion strategies to promote their propagation, which includes host-translational shut-off (30). Indeed, while HCV infection results in effective IRG induction, virus induced phosphorylation of PKR inhibits eIF2α and therefore blocks host IRG protein translation (31). PKR activation is therefore advantageous to HCV and prevents clearance because while translation of capped host mRNAs are dependent on eukaryotic initiation factors (eIFs), HCV polyprotein translation occurs independently of eIFs using an IRES located in the 5’UTR of the viral genome (32). Further to the described HCV inhibition of eIF2α (31), our transcriptional profiling identifies broad down regulation of multiple components of the host translational machinery. Mechanistically, this process is likely mediated by the TF MYC, which is known to directly regulate ribosome biogenesis and translation, controlling the expression of RPS and RPL proteins of the small and large ribosomal subunits, in addition to the gene products necessary for rRNA processing, nuclear export of ribosomal subunits and mRNA translation initiation (33). Interestingly, in PHHs, significant upregulation of *MYC* mRNA results in down-regulation of the gene products it controls (Fig 6C, right panels). These data may represent a previously unappreciated mechanism whereby downregulation of constituent components of the translational machinery arrests the translation of antiviral effector genes. Contrastingly, MYC is predicted to be inhibited in upstream regulator analysis (Fig. 5A). Detailed investigation of the genes involved in the prediction MYC inhibition and the interactions included in the IPA knowledge base highlights a number of genes involved in IFN-signaling based on microarray analysis in plasmacytoid dendritic cells (pDC) (34). These genes are highly upregulated upon *MYC* knockdown in pDCs and the same genes are induced in the antiviral immune response in PHH upon HCV infection. Differences between cellular responses in pDCs and PHH or upon knockdown of MYC compared to HCV infection may result in upregulation of *MYC* in our data set. Together, these data simultaneously capture infection-induced transcriptional signatures associated with pro-viral translational shut-off and anti-viral IFN signaling.

Furthermore, we observed transcriptional dysregulation of additional gene programs which have been reported to modulate susceptibility to HCV. We observed targeting of the mTOR signaling pathway, which has been reported as pro-viral, with mTOR inhibitor rapamycin targeting HCV replication in vitro and reducing viral RNA levels in patients post transplantation (35). HCV infection has also been shown to activate mTOR and pharmacological inhibition of mTOR was shown to suppress HCV virion assembly and release in vitro (36). Of note, significant targeting of multiple pathways associated with nuclear receptor (NR) signaling was detected in both PHHs and Huh-7.5 cells, further confirming that Huh-7.5 cells retain some HCV-inducible programs that are of biological relevance. NR-mediated signaling regulates transcriptional programs that control host metabolic processes and lipid metabolism and are reported to modulate susceptibility to HCV infection in both humans (37) and mice (38).

In addition to HCV activation of pro-and antiviral cascades, we also investigated the contribution of *IRF1*-mediated intrinsic immunity in PHHs to the control of HCV replication. IRF1 is a TF that participates in IFN induction but also directly induces a subset of IRGs. Additionally, *IRF1* represents a potent pan-viral restriction factor and a key component of the cellular antiviral response (7). Using microarray analysis, Schoggins *et al*. identified a panel of 130 partially overlapping, *IRF1*-regulated genes (>3-fold) via lentiviral over-expression in Huh-7 and *STAT1*^-/-^ fibroblasts. More recently it has been demonstrated that constitutive expression of *IRF1* in immortalized PH5CH5 cells of hepatic origin (24) and BEAS-2B bronchial epithelial cells (23) coordinates intrinsic antiviral protection independently from the IFN system. Ectopic expression of *IRF1* in Huh-7.5 cells enabled us to define a greatly expanded *IRF1* regulon, and comparative statistical analysis of Huh-7.5 EMPTY, Huh-7.5 *IRF1* and uninfected PHHs transcriptomes provides supportive evidence that baseline immunity in PHHs in the absence of infection is orchestrated by *IRF1*. Previous studies have shown *IRF1* to be a potent restrictor of HCV replication (7). We confirm this observation and also determine that *IRF1* can significantly inhibit HCV subgenome translation in Huh-7.5 cells. While intrinsic *IRF1* expression in PHHs maintains a suite of genes that can actively suppress HCV translation and replication, which may contribute to the low-levels of gene induction at 6 hpi, this translational suppression could be partially overcome by the host translational shut-off we observe at 72 hpi.

In summary, the virus-host interactions which determine the susceptibility of human hepatocytes to initial HCV infection and their capacity to support persistent infection are incompletely defined. The high levels of genetic diversity seen between HCV genotypes point to a long period association for virus and host co-evolution (39) and HCV has evolved multiple strategies to hijack the host cell machinery required to facilitate it propagation, while at the same time evading host defenses. We observe early concurrent transcriptional dysregulation of gene programs which facilitate viral persistence, balanced against those which promote viral inhibition. This cellular pro-and anti-viral antagonism may keep HCV replication levels below a clearance threshold in the initial phase of infection and ultimately facilitate progression to chronicity.

## MATERIALS AND METHODS

### Source of PHHs

PHHs were isolated from surgical liver resections as previously described (18). PHHs were obtained with informed consent approved by the ethics commission of Hannover Medical School (Ethik-Kommission der MHH, #252-2008). Additionally, PHHs were commercially obtained (Lonza, Basel, Switzerland). Cryopreserved PHH were thawed as recommended by the manufacturer.

### Virus production and UV-inactivation

Huh-7.5 cells were electroporated with HCV genomic RNA transcripts (Jc1 strain) (40, 41) and supernatants were harvested at 48, 72 and 96 hours post-electroporation, filtered (0.45 micron pores), pooled, aliquoted and frozen at -80°C. A portion of this virus stock was UV-inactivated in 6-well-dish with 1ml per well at 5J/cm^2^.

### Generation of Huh-7.5 *IRF1* cells

The *IRF1*-coding sequence (gBlock®, IDT) was cloned into lentiviral vector pWPI-bla (Addgene) and confirmed by Sanger sequencing (GATC). Co-transfection of plasmids encoding VSV-G, HIV-1gag/pol and pWPI-bla-*IRF1* into HEK293T cells was performed using Lipofectamine 2000 (Invitrogen). Supernatants containing lentiviral pseudoparticles for transgene delivery were harvested at 24 h and 48h, pooled, filtered (0.45 micron pores) and used to transduce 2×10^5^ Huh-7.5 cells. Seventy-two hours post transduction, blasticidin (10 ug/ml) was added to media. For control purposes, Huh-7.5 cells were also transduced with pseudoparticles containing an empty pWPI-bla vector (no transgene expressed). After blasticidin addition, surviving cells were expanded via passaging for 14 days prior to freezing at -150°C with 10% DMSO.

### HCV infection and RNA isolation

PHH infections were performed at a multiplicity of infection of 1 (MOI 1) calculated on Huh-7.5 cells and the same volume was used for treatment with UV-inactivated virus. Four hours post-inoculation, cells were washed with PBS and 1 ml of fresh Hepatocyte Culture Medium (HCM) was added per well (Lonza). At 6 or 72 hpi, supernatants were collected and cells were lysed in 1 ml TRIzol® reagent (Invitrogen) or RA1 buffer supplemented with β-mercaptoethanol (Macherey-Nagel). All samples were stored at -80°C until processing. Total RNA was extracted according to the manufacturer’s instructions. For infection of Huh-7.5 empty vector transduced cells, identical conditions were used except DMEM was used instead of HCM media and RNA extractions were performed using only the NucleoSpin RNA kit (Macherey-Nagel). Where ruxolitinib treatment is indicated, PHHs were pretreated with 10µM ruxolitinib on the day before HCV infection. For the rescue of viral particle production (Fig 2C), 10µM ruxolitinib was also added to HCM media directly after infections.

### Subgenomic replicon assays

Huh-7.5 cells expressing *IRF1* or transduced with an EMPTY lentivirus were electroporated with equal amounts (2.5µg) of a firefly luciferase (F-luc) expressing subgenomic replicon RNA (NS3-NS5B, strain JFH-1) and seeded onto 12-well dishes. Additionally, EMPTY control cells were treated with 1 µM 2’CMA or DMSO. Cells were lysed at 4, 24, 48, 72 and 96 hours post electroporation in 350 µl passive lysis buffer (Promega) per well and frozen at -20°C until measurement of F-luc expression with a tube luminometer.

### TCID_50_ and RT-qPCR

Viral titers in cellular supernatants and viral stocks used for infection experiments were quantified on Huh-7.5 cells using a limiting dilution assay as previously described (42). The limit of quantification was determined by the lowest, still evaluable result in the assay set up. A quantitative, one-step RT-qPCR was performed to determine intracellular HCV-RNA copy numbers, using the “Light Cycler 480 RNA Master Hydrolysis Probes” kit (Roche) and an HCV specific primer-probe set targeting the 5’ UTR region. RT-qPCR was performed using a Light Cycler 480 (Roche). Ten RNA copies was the lowest number used to calculate the standard curve.

To determine relative gene expression of selected cellular genes, 250 ng of total cellular RNA were reverse transcribed using the Takara Reverse Transcription (RT) kit. Takara’s SYBR Premix Ex Taq II was used according to manufacturers’ instructions with gene specific primers. RT-qPCR was performed using a Light Cycler 480 (Roche). Changes in relative gene expression were calculated according to the 2^-ΔΔCT^ method (43).

### RNA-seq

Cellular RNAs were used to generate sequencing libraries using a ScriptSeqv2 kit (Illumina), run on the Illumina HiSeq 2500 platform, and subsequent data analyses were performed using CLC Genomics Workbench (Qiagen, Aarhaus) (44). Mapping against human reference genome (hg38) was performed for individual samples and relative transcript expression was calculated from raw count data via normalization to gene length (RPKM) (45). Identification of differentially expressed genes (DEGs) was conducted by comparing raw count data, with calculation of false discovery rate (FDR) p-values for multiple comparisons. Significant DEGs with low expression (FDR p-values <0.05 with a final RPKM <1) were omitted from subsequent GO, TF and pathway analyses.

### GO enrichment analyses

GO analyses were performed using the GO Resource (http://geneontology.org/). ENSEMBL identifiers for DEGs were used as input and identification of significantly enriched GO categories was performed using the Panther Classification system (http://pantherdb.org/). *P*-values for specific GO categories were generated after Bonferroni correction for multiple testing. Redundant GO terms were removed with REVIGO (46) (http://revigo.irb.hr) with the following settings: allowed similarity “tiny”, GO term sizes database “Homo sapiens” and “SimRel” semantic similarity measure. Remaining terms were visualized in semantic similarity-based scatterplots using GraphPad Prism v9.0.

### Canonical pathway analysis and transcriptional regulators

Pathway analyses were performed using Ingenuity Pathway Analysis (Qiagen, Aarhaus) (47). Input data included ENSEMBL identifiers, FDR-p-values and fold change values for DEGs determined using CLC genomics workbench. Default settings were used for all categories except for species, which was set to human. Z-scores were calculated based on the expression fold change. IPA compares input data sets with the ingenuity knowledge base which represents a collection of published interactions of molecules. Upstream regulators are determined bioinformatically (http://pages.ingenuity.com/rs/ingenuity/images/0812%20upstream_regulator_analysis_whitepaper.pdf). Analyses outputs were exported as excel sheets or pdf files and data re-plotted with GraphPad and Adobe illustrator. Disease-, cancer-and non-hepatocyte-associated pathways were omitted from presented diagrams

## Data Availability

RNA-seq data generated in this study and subsequent downstream analyses including identification of DEGs, enriched GOs categories, upstream TFs and targeted pathways are submitted to the NCBI GEO database (GEO accession number GSE166428).

## ACKNOWLEDGEMENTS

We thank Charles Rice (Rockefeller University) for Huh-7.5 cells and anti-NS5A antibody 9E10. We also thank Robert Geffers, Michael Jarek, Maren Scharfe and Sabin Bhuju (HZI Braunschweig, GMAK facility) for RNA-seq library preparation and generation of HiSeq 2500 raw data. RJPB was supported by BMG grant 1-2516-FSB-416. TP was supported by a grant from the European Research Council ERC-2011-StG_281473-(VIRAFRONT). TP was also funded by the Deutsche Forschungsgemeinschaft (DFG, German Research Foundation) under Germany’s Excellence Strategy –EXC 2155 “RESIST”-project ID 39087428.

